# Reemergence of yellow fever virus in southeastern Brazil, 2017-2018: what sparked the spread?

**DOI:** 10.1101/2021.09.02.458267

**Authors:** Joelle I. Rosser, Karin Nielsen-Saines, Eduardo Saad, Trevon Fuller

## Abstract

**Background:** The 2017-2018 yellow fever virus (YFV) outbreak in southeastern Brazil marked a reemergence of YFV in urban states that had been YFV free for nearly a century. Unlike earlier urban YFV transmission, this epidemic was also driven by forest mosquitos. The objective of this study was to evaluate environmental drivers of this outbreak.

**Methodology/Principal Findings:** Using surveillance data from the Brazilian Ministry of Health of human and non-human primate (NHP) cases of yellow fever, we traced the spatiotemporal progression of the outbreak. We then assessed the epidemic timing in relation to drought using a monthly Standardized Precipitation Evapotranspiration Index (SPEI). Lastly, we evaluated demographic risk factors for rural or outdoor exposure amongst YFV cases. Both human and NHP cases were first identified in a hot, dry, rural area in northern Minas Gerais before spreading southeast into the more cool, wet urban states of Espírito Santo, São Paulo, and Rio de Janeiro. Outbreaks also coincided with drought in all four southeastern states of Brazil. Confirmed YFV cases had an increased odds of being male (OR 2.58; 95% CI 2.28-2.92), working age (OR: 2.03; 95% CI: 1.76-2.35), and reporting recent travel from an urban to a rural area (OR: 5.02; 95% CI: 3.76-6.69).

**Conclusions/Significance:** The 2017-2018 YFV epidemic in Brazil originated in hot, dry rural areas of Minas Gerais before expanding south into urban centers. An unusually severe drought in this region may have created environmental pressures that sparked the reemergence of YFV in Brazil’s southeastern cities.

**Author Summary:** In 2017-2018, cities in southeastern Brazil experienced an unusual outbreak of yellow fever virus. In the early 20^th^ century, these cities had large outbreaks of yellow fever, spread by *Aedes* mosquitos. But until this recent outbreak, they had been free of yellow fever for nearly a century. While this outbreak was spread by *Haemagogous* forest mosquitos, the reemergence of yellow fever in densely populated urban areas raises serious concerns about it reestablishing ongoing transmission in cities, spread by urban *Aedes* mosquitos. Our study sought to understand how and why yellow fever virus remerged in this area. We traced the outbreak, finding that it started in hot, dry, rural areas and spread south into cool, wet urban areas. Additionally, the outbreak coincided with a severe drought; this extreme weather may have promoted the spread of yellow fever. Infection was also associated with rural and outdoor exposure, further suggesting this epidemic originated in rural areas.

## Introduction

Between December 2016 and July 2019, a YFV outbreak occurred in southeastern Brazil, resulting in over 2,000 confirmed human cases[1,2]. The height of this epidemic occurred during the summer seasons of 2017 and 2018. This outbreak was unusual both in its large size and geographic distribution. Urban transmission cycles involving *Aedes sp*. mosquitos were responsible for severe YFV epidemics up until the early 20^th^ century when aggressive vector-control and vaccination campaigns eliminated this type of transmission pathway[3]. However, since the early 1940’s, YFV in Brazil has circulated exclusively by a sylvatic transmission cycle in which *Haemagogus sp*. mosquitos, which live and breed in forest canopies, feed primarily on non-human primates (NHPs) and only sporadically feed on humans[4–6]. This results in low level transmission with periodic small outbreaks occurring in rural northern and western forest states of Brazil.

In contrast, the 2017-2018 epidemic occurred in southeastern Brazil, including São Paulo, Espírito Santo, and Rio de Janeiro, urban states which had not experienced YFV transmission in nearly a century. Outbreaks of YFV in these urban states raise great concern for the possibility of an urban transmission cycle resurgence involving *Aedes aegypti* and *Aedes albopictus*[7,8]. These two mosquitos preferentially breed in small water containers and therefore thrive in urban slums with poor water infrastructure[9,10]. In recent decades, Espírito Santo and Rio de Janeiro have experienced several large outbreaks of other flaviviruses such as dengue, chikungunya, and Zika which are transmitted by these *Aedes* mosquitos[11–13]. If YFV reestablished an urban cycle involving *Aedes* in these densely populated, inadequately vaccinated states, the public health impact could be enormous.

Curiously, albeit fortunately, there has been no evidence of *A. aegypti* infection in the 2017-2018 YFV epidemic[8,14]. This in no way precludes the possibility of a future spillover to *Aedes aegypti*. However, this particular epidemic was driven by conditions promoting transmission by *Haemagogus* mosquitos[14]. The recent incursion of YFV in the southeastern states of Brazil suggests that a change in environmental conditions triggered a new transmission dynamic in southeastern Brazil. While the 2017-2018 YFV epidemic included three previously unaffected states, it started in the neighboring state of Minas Gerais which did start having low level YFV transmission in previous years[1]. Minas Gerais is a rural state uniquely positioned in a transition zone between forested areas to the northwest and the southeastern coastal states[15]. It is however distinct from Amazon-like forested areas and is comprised predominantly of a savannah-like *cerrado* biome on the western side of the state and the Atlantic rainforest on the eastern side (“Mata Atlântica”) which is contiguous with São Paulo, Espírito Santo, and Rio de Janiero [2,16].

This study aims to evaluate environmental and demographic predictors of the YFV outbreak in southeastern Brazil. We hypothesize that a severe drought during this time set the stage for YFV transmission to intensify at the rural-urban interface and then spread into urban areas. To assess this hypothesis, we traced the evolution of human and non-human primate (NHP) cases over space and time to demonstrate how the epidemic spread from rural, hot, dry parts of the southeastern region of Brazil to more urban, cool, wet areas. We also evaluated the timing of the outbreaks in relationship to a drought index. Finally, we evaluated demographic characteristics of human YFV cases to assess risk factors for rural and outdoor exposures.

## Methods

### Geographic distribution of human and NHP cases over time

The number of confirmed human YFV cases by municipality were obtained from the Sistema de Informação de Agravos de Notificação (SINAN)[17]. Suspected cases of YFV were classified as confirmed in the SINAN database based on whether the case had serologic or polymerase chain reaction evidence of an acute yellow fever virus infection. Southeastern Brazil experiences two main seasons: the more cool and dry ‘winter’ season from May to October and the more hot and wet ‘summer’ season from November to April, with peak rainfall and high temperatures in January to February. Based on reported date of symptom onset and seasonal changes, cases were divided into time periods to show the progression of case counts over the course of the epidemic.

We also obtained data from SINAN on the number of non-human primates (NHPs) that were found dead, tested positive for YFV, and were reported by state health departments to the Division of Zoonoses and Vector-borne Diseases of the Ministry of Health from January 2007 to December 2020 (data received from SINAN)[17]. The surveillance included five genera of NHPs: Alouatta, known as howler monkeys which are small nocturnal frugivores, Callithrix, which are small marmosets whose diet consists of insects, fruits and other plants, and Cebus, Saimiri, and Sapajus, all of which are medium-sized NHPs that live in large troops [2]. We pooled the genera and created a map of the municipalities reporting at least one confirmed NHP case in each of 2016, 2017, and 2018.

Human and NHP cases were mapped to municipality in R Studio version 1.1.456 using the geobr package[18] which pulls municipality boundaries from the Instituto Brasileiro de Geografia e Estatistica (IBGE)[19].

### Geographic distribution of temperature, rainfall, and population density across southeastern states of Brazil

Raster data for the historical (1970-2000) monthly mean, minimum, and maximum temperature and mean rainfall was obtained from WorldClim[20] and then mapped for the southeastern states of Brazil to depict averages during the summer season (November to April) and winter season (May to October) over this geographic region using R Studio and the geobr package[18]. Data on population density in 2017 across this region was obtained from WorldPop and mapped in R Studio [21].

### Drought

To assess drought conditions, we obtained precipitation and temperature for each month from January 2007 to December 2020 from weather stations in each state. We queried the databases of the Brazilian National Institute of Meterology and National Water Agency for all stations in each state and selected the weather station nearest to the municipality with the greatest number of human cases of YFV. If two stations were equidistant from the municipality, we selected the one with the fewest missing observations during the study period. This resulted in the selection of weather stations in Vitória, Espírito Santo, São Paulo, São Paulo, Teófilo Otoni, Minas Gerais, and Campos dos Goytacazes, Rio de Janeiro.

These data were used to calculate the monthly Standardized Precipitation Evapotranspiration Index (SPEI), which is a measure of water balance based on rainfall and temperature [22]. A negative value of the index indicates that the evapotranspiration exceeded precipitation, resulting in a water deficit. Months in which the SPEI assumed a negative value were anomalously hot and dry, compared to the average climatic conditions during the study period. We defined drought as months in which the SPEI was less than zero. To calculate the index, we utilized the aforementioned weather station data and the R package SPEI 1.7. The SPEI settings consisted of a scale parameter of 8 months and a Gaussian kernel.

### Rural /outdoor exposure

Rural or outdoor exposure was assessed based on relevant demographic characteristics including sex, working age, recent travel history, and occupation as reported to SINAN in 2017 and 2018[17]. Working age was defined as being between the ages of 16 and 65 years old, the minimum legal working age and the retirement age for men in Brazil, respectively. Travel history was divided into five categories based on the direction of travel between rural and urban areas. Age, gender, and travel history were compared between suspected cases of YFV which were confirmed versus those that were discarded using Fisher’s exact test.

For the subset of confirmed cases for whom occupation was recorded, occupation was classified into one of four categories: farmer / rural worker, other outdoor worker, indoor worker, or ‘intermediate’ (for jobs which could be both indoors and outdoors). To assess whether rural/outdoor occupational risk was higher at the beginning of the epidemic, the number of confirmed cases with a known occupation of farmer/rural worker versus other occupations were compared between the first (2017) and second (2018) years of the epidemic and between the start of each new wave (January) and subsequent months (February through December) using Fisher’s exact test.

The human data used in this analysis was deidentified data available from an existing public access database (SINAN) so human subjects ethical review was not required.

## Results

### Geographic distribution of human and NHP cases over time

The first human cases of YFV in 2016 were identified in December in the northeastern region of Minas Gerais. This geographic area is one of the hottest and driest parts of the southeastern region of Brazil. In January 2017, case counts increased in intensity and spread outwards from that initial epicenter towards the border of Minas Gerais and Espírito Santo, still a relatively hot and dry region. For the remainder of the 2017 summer season (ending in May) the epidemic spread predominately through Espírito Santo, with smaller numbers of cases in the cooler, wetter states of Rio de Janeiro and São Paulo. A few scattered cases were identified in all four states during the winter season. When the epidemic reemerged during the following January 2018, it centered around the cities in the more southeastern part of this region, including São Paulo, Rio de Janeiro, and Belo Horizonte. (Figure 1)

**Figure 1.**
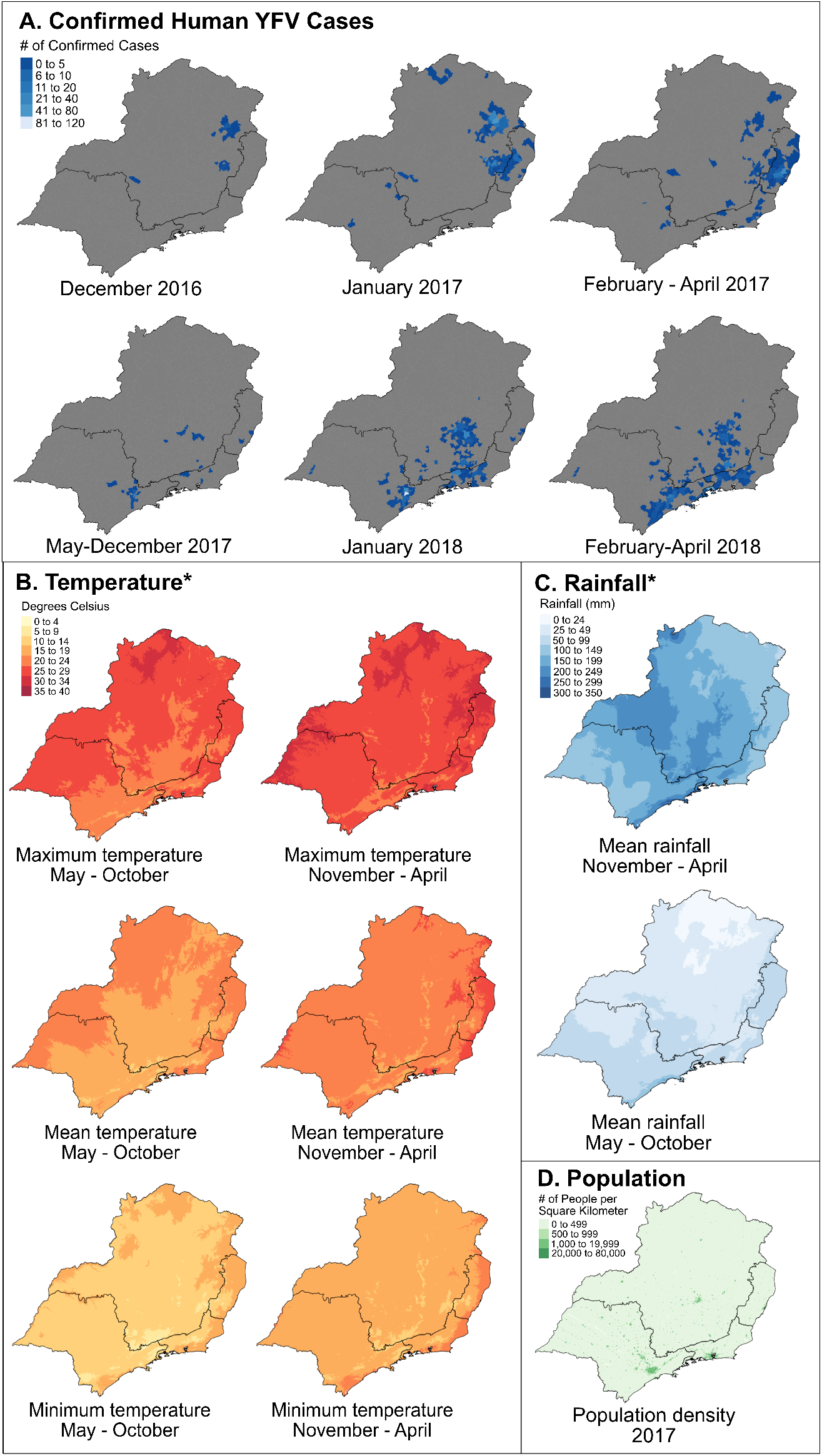
Yellow fever virus outbreak emergence in hot, dry rural areas of southeastern Brazil. Abbreviations: YFV = yellow fever virus; mm = millimeters. *Temperature and rainfall values are derived from the historical average (1970-2000) from WorldClim[20]. A. The YFV outbreak was first identified in the northeastern region of Minas Gerais and subsequently spread east into Espírito Santo and then south into São Paulo and Rio de Janeiro. B. The northeastern region of Minas Gerais and Espírito Santo experiences the highest temperatures in this region. Maps on the left show the cooler winter months and on the right show the warmer summer months. C. The northern area of Minas Gerais is also the driest part of this region. D. The northeastern region of Minas Gerais is also the most rural and least densely populated part of this region.

The geospatial progression of human cases roughly paralleled that of NHP cases over this time (Figure 2). Scattered NHP cases were identified in rural Minas Gerais and São Paulo throughout 2016. There is one area in the northernmost part of Minas Gerais where multiple municipalities had NHP cases in 2016. There are several municipalities bordering this area which had NHP cases in both 2016 and 2017 or 2017 alone. This apparent progression of NHP cases extended southeast from this rural northern region towards the early epicenter (in December 2016/January 2017) of human YFV cases in northeastern Minas Gerais. By 2018, the NHP cases were predominantly centered around municipalities in the more urban, southeastern part of this region, again overlapping with the human epidemic. (Figure 2)

**Figure 2:**
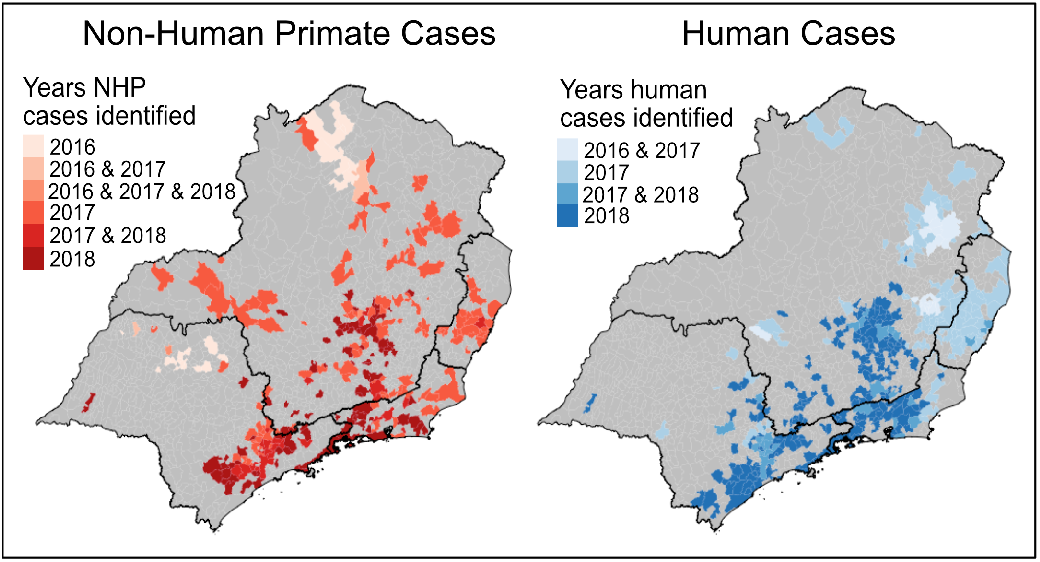
Human & NHP Yellow Fever Viruses Cases by Year. Abbreviations: NHP = Non-human primate. NHP cases (left map) were identified in the rural, northern area of Minas Gerais starting in 2016 and continuing through the epidemic in 2017 and 2018. The earliest human cases identified during this outbreak also occurred in northern Minas Gerais in 2016 and 2017. As the epidemic progressed through 2017 and 2018, both NHP and human cases spread south into the urban centers of Rio de Janeiro and São Paulo.

### Drought

A major drought hit southeastern Brazil starting around 2012 and ending around 2019. We plotted a drought index as measured by the Standardized Precipitation Evaporation Index (SPEI) from January 1, 2007 to December 31, 2020 against confirmed yellow fever cases in each of the four southeastern states of Brazil (Figure 3). The model fit between monthly cases and monthly SPEI as measured by Pearson’s correlation coefficient for Minas Gerais, Espírito Santo, Rio de Janiero, and São Paulo were 0.17, 0.16, 0.11, and 0.28, respectively. When limiting this analysis to peak summer months (December to April), when YFV transmission occurs, Pearson’s correlation coefficients mildly increased to 0.26, 0.27, 0.14, and 0.16. This relatively low model fit is driven largely by the several months of drought in which there were no cases of YFV.

**Figure 3.**
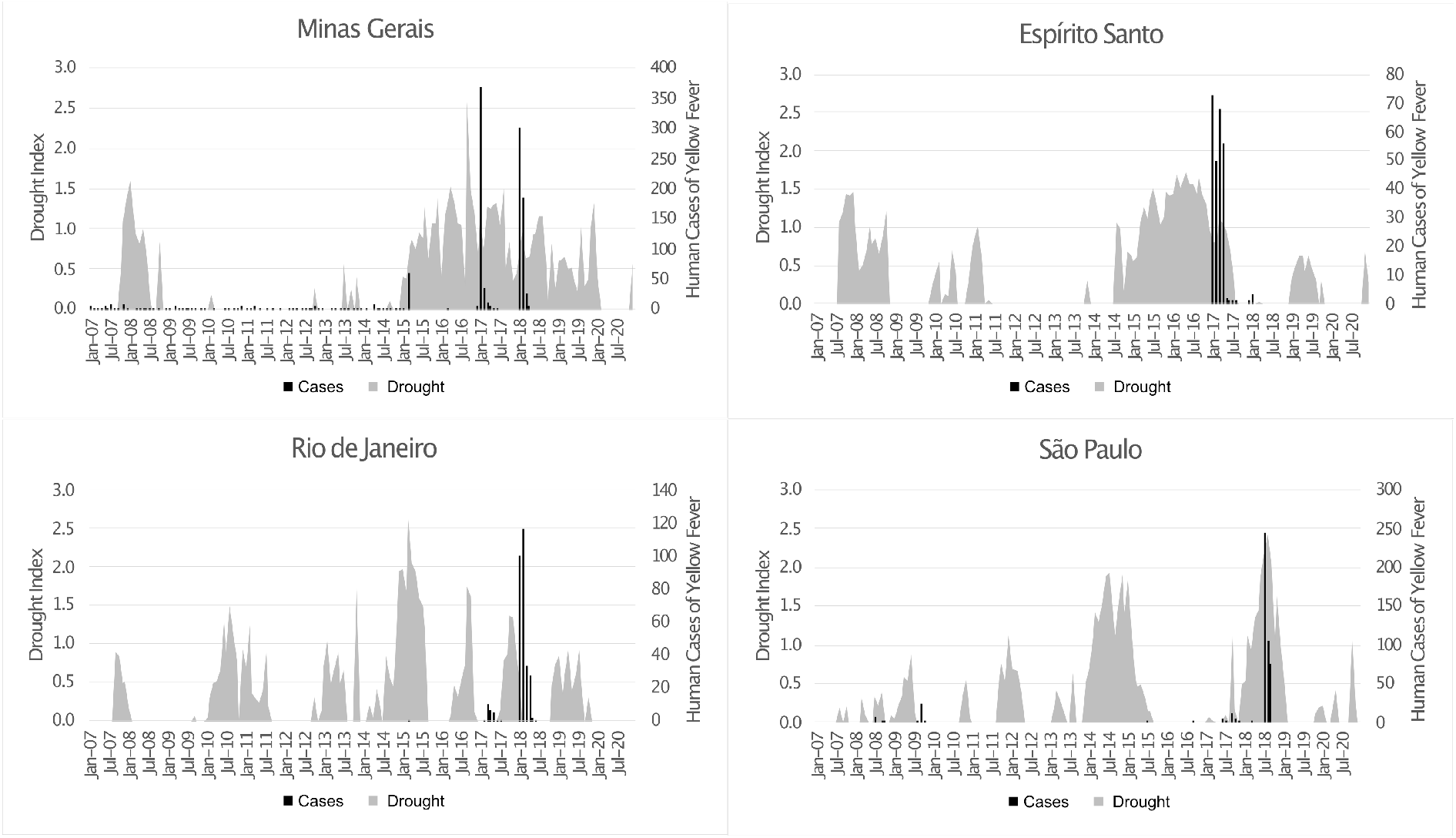
Timing of Drought & Yellow Fever Outbreaks. Abbreviations: Jan = January; Jul = July. The “Drought Index” is derived from the Standardized Precipitation Evapotranspiration Index (SPEI) which measures water balance based on rainfall and temperature. A negative value of SPEI indicates a water deficit; a negative SPEI is represented here by grey bars showing an increased drought index. The number of yellow fever virus cases reported in each state is represented by the black bars. Yellow fever virus outbreaks coincided with severe drought conditions in each of the four southeastern states of Brazil.

However, the peak of the YFV epidemic did coincide with an elevated drought index in all four southeastern states. The drought index, SPEI, was consistently over 1.0 (moderate drought) for a period of two to three years in both Minas Gerais and Espírito Santo when the epidemic hit those states in 2016, and in Minas Gerais the SPEI exceeded 2.0 (extreme drought) at the end of 2016 just prior to the start of the epidemic in that state. In Minas Gerais, the SPEI remained high in 2018 and a second wave of the epidemic hit in 2018, whereas in Espírito Santo, the SPEI returned to zero shortly after the 2017 epidemic and this state was almost entirely spared from a second wave of cases in 2018.

Rio de Janeiro and São Paulo, which were most affected by the second wave of the epidemic in 2018, had shorter and more intermittent spikes in the drought index in the years leading up to the epidemic, rather than the multiple years of sustained elevated SPEI experienced by the more northern states of Minas Gerais and Espírito Santo. Notably, the SPEI in 2016 and the first half of 2017 (during the first wave of the epidemic further north) for both Rio de Janiero and São Paulo was near zero and then increased to nearly 2.0 (extreme drought) in early 2018, again coinciding with the peak of the epidemic in those states.

### Rural /outdoor exposure

Rural and outdoor exposure was assessed by demographic characteristics including sex, age, recent travel history, and occupation. Confirmed cases of YFV were significantly more likely to be male (Odds Ratio (OR): 2.58; 95% Confidence Interval (CI): 2.28-2.92) and be of working age (OR: 2.03; 95% CI: 1.76-2.35) compared to discarded cases (Table 1). A recent history of travel was also significantly associated with confirmed cases of YFV. Travel from an urban area to a rural area was the highest risk travel category (OR: 5.02; 95% CI: 3.76-6.69). The inverse, travel from a rural area to an urban area, was the only travel category not significantly associated with an increased risk of confirmed YFV infection.

**Table 1.**
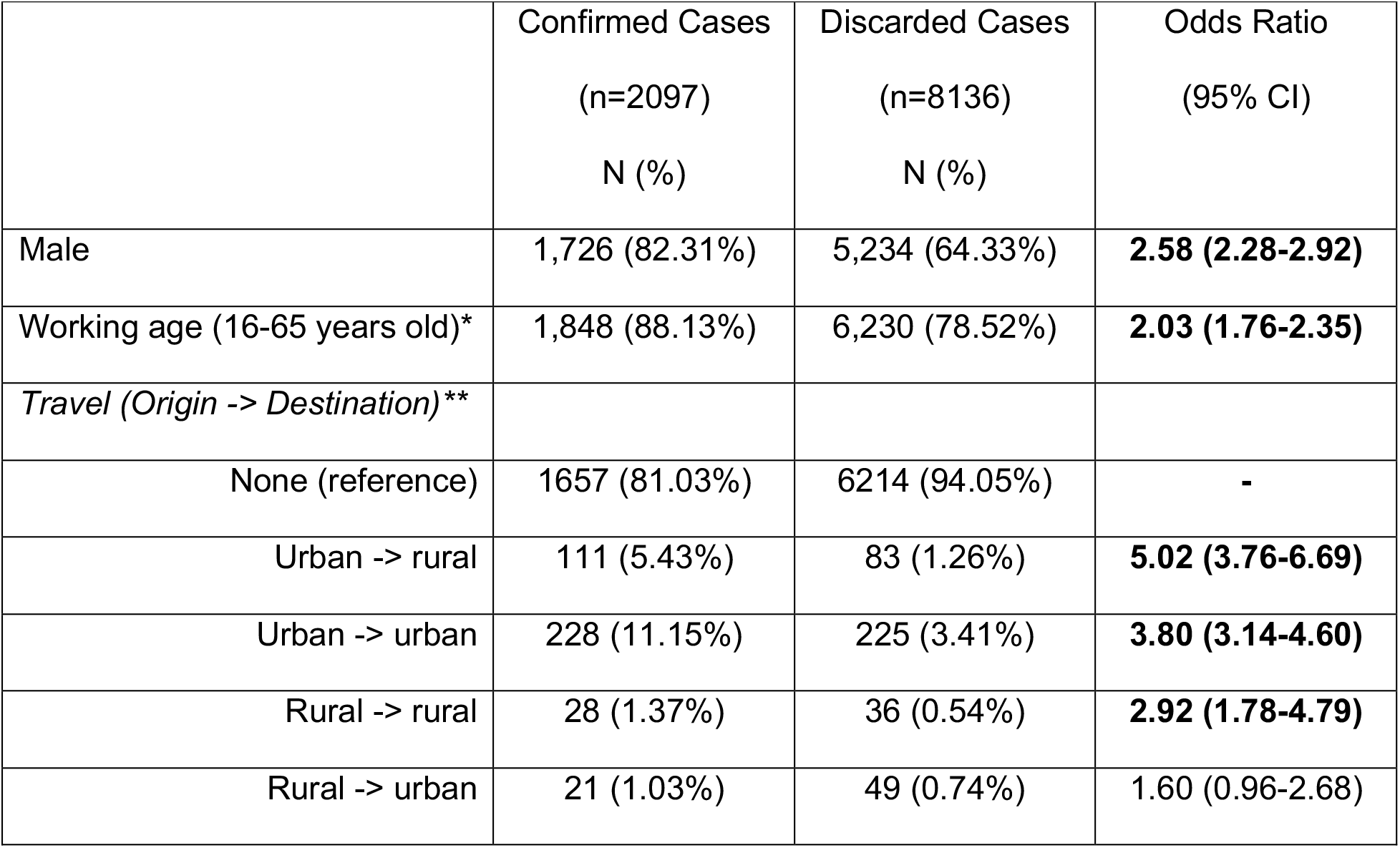
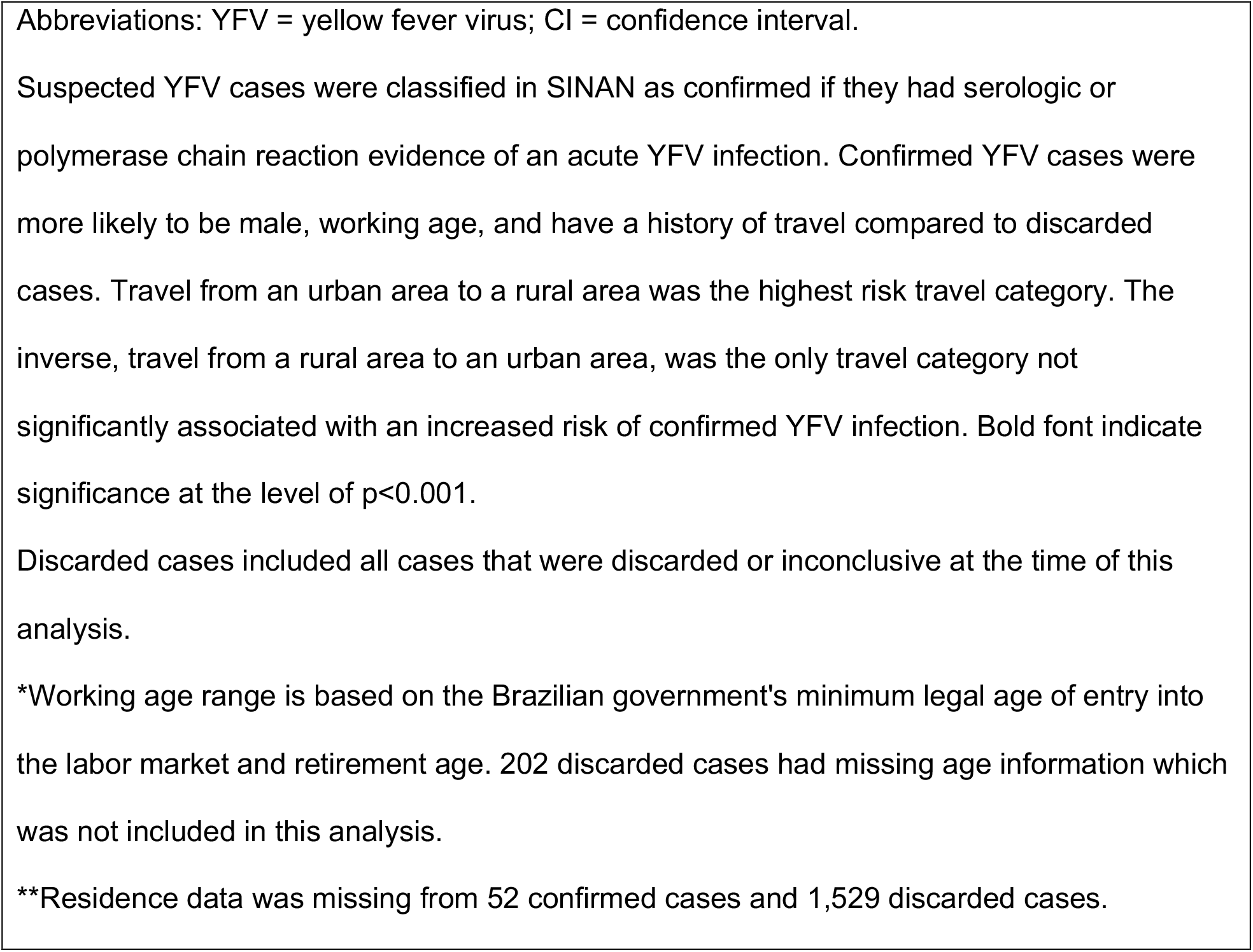
Demographic characteristics of confirmed versus discarded YFV cases in 2017 & 2018.

|  |  |  |  |
| --- | --- | --- | --- |
|  | Confirmed Cases<br>(n=2097)<br>N (%) | Discarded Cases<br>(n=8136)<br>N (%) | Odds Ratio<br>(95% CI) |
| Male | 1,726 (82.31%) | 5,234 (64.33%) | **2.58 (2.28-2.92)** |
| Working age (16-65 years old)* | 1,848 (88.13%) | 6,230 (78.52%) | **2.03 (1.76-2.35)** |
| *Travel (Origin -> Destination)\*\** |  |  |  |
| None (reference) | 1657 (81.03%) | 6214 (94.05%) | - |
| Urban -> rural | 111 (5.43%) | 83 (1.26%) | **5.02 (3.76-6.69)** |
| Urban -> urban | 228 (11.15%) | 225 (3.41%) | **3.80 (3.14-4.60)** |
| Rural -> rural | 28 (1.37%) | 36 (0.54%) | **2.92 (1.78-4.79)** |
| Rural -> urban | 21 (1.03%) | 49 (0.74%) | 1.60 (0.96-2.68) |

Abbreviations: YFV = yellow fever virus; CI = confidence interval.
Suspected YFV cases were classified in SINAN as confirmed if they had serologic or polymerase chain reaction evidence of an acute YFV infection. Confirmed YFV cases were more likely to be male, working age, and have a history of travel compared to discarded cases. Travel from an urban area to a rural area was the highest risk travel category. The inverse, travel from a rural area to an urban area, was the only travel category not significantly associated with an increased risk of confirmed YFV infection. Bold font indicate significance at the level of $p < 0.001$ .
Discarded cases included all cases that were discarded or inconclusive at the time of this analysis.
\*Working age range is based on the Brazilian government's minimum legal age of entry into the labor market and retirement age. 202 discarded cases had missing age information which was not included in this analysis.
\*\*Residence data was missing from 52 confirmed cases and 1,529 discarded cases.

Occupation was available for 765 (36.5%) of the 2,097 confirmed cases in 2017 and 2018. Farmers/rural workers made up the largest occupational category with 398 (52.0%) of the 766 cases. Confirmed cases were more likely to be a farmer/rural worker than another occupation in 2017 compared to 2018 (OR: 3.16; 95%CI: 2.32-4.32) and in January compared to other months (OR: 1.81; 95%CI: 1.34-2.45). (Figure 4)

**Figure 4.**
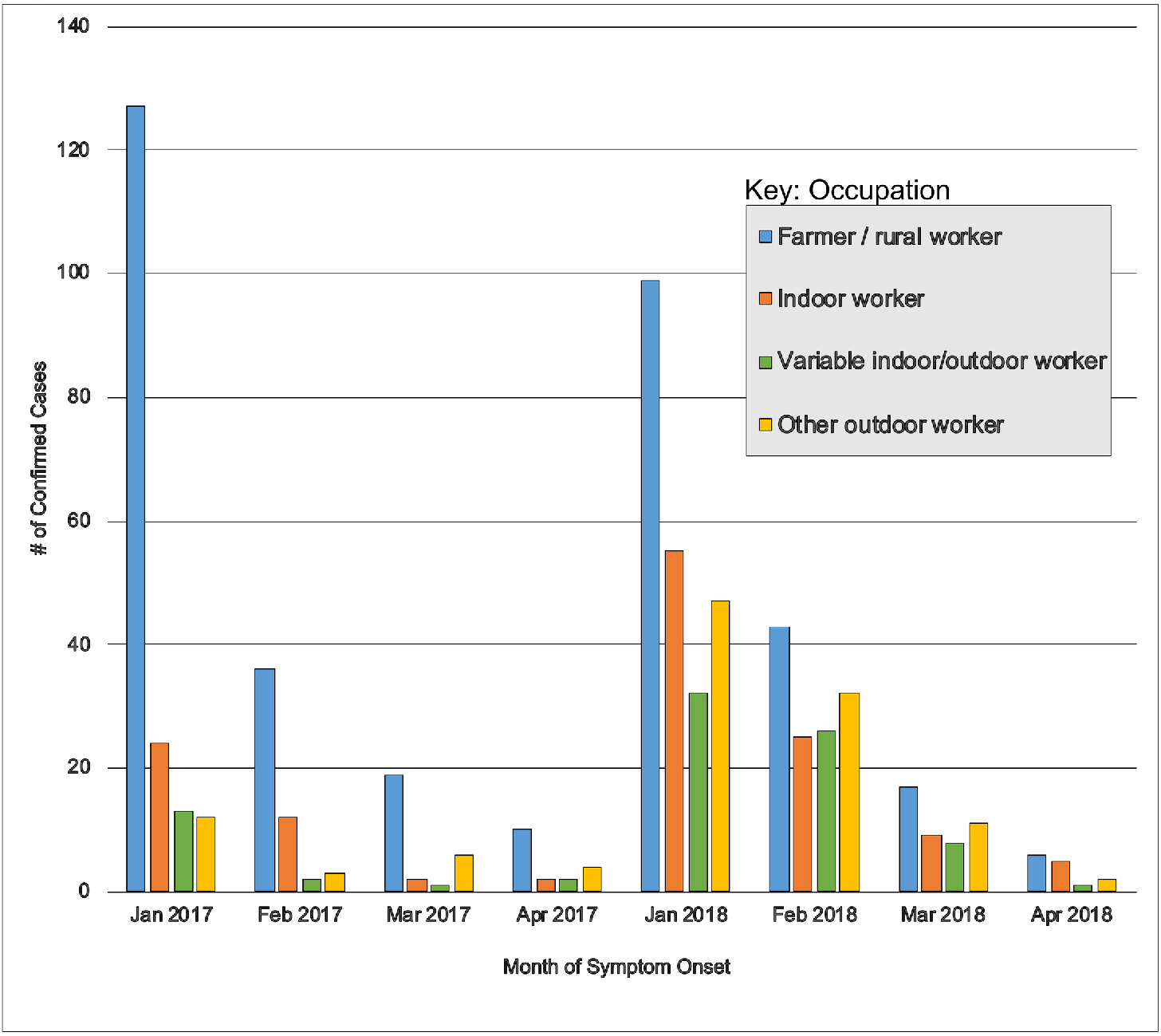
Occupation of yellow fever virus cases over the course of the epidemic. Abbreviations: Jan = January; Feb = February; Mar = March; Apr = April. Farmers and rural workers (blue bars) dominated the cases in the first year of the epidemic (2017). This was also the predominant group at the start of each wave of the epidemic (January 2017 & January 2018).

## Discussion

Our results show that the YFV outbreaks of 2017-2018 coincided with periods of drought in all four southeastern states of Brazil affected by this unusual epidemic. We also trace the epidemic as it starts in the hot, dry northern region of Minas Gerais and spreads east and south into cooler, damper, and more urbanized areas. The pattern of this spatiotemporal progression is seen with both confirmed human and NHP cases. Lastly, we found an increased risk of yellow fever associated with demographic characteristics suggestive of outdoor and rural exposure, including male sex, working age, and recent travel particularly from an urban to a rural area; additionally, farmer/rural worker was the most common work category, particularly at the start of outbreaks. These findings are consistent with small genomic sequencing studies that suggest multiple introductions of YFV from Minas Gerais into the more urban states of Espírito Santo, Rio de Janeiro, and São Paulo [23–25]. This supports our hypothesis that this epidemic started in rural areas of Minas Gerais where intermittent transmission had previously occurred and that environmental conditions during this time were uniquely primed for YFV spread to neighboring states and urban centers where huge outbreaks occurred. This is also consistent with other studies that have demonstrated an overall southeastern spread of yellow fever through Brazil over the past two decades and during this epidemic in particular[26].

This 2017-2018 YFV outbreak occurred in a region of Brazil that had not experienced urban YFV transmission in nearly a century. Around the same time, Brazil experienced one of the most extreme droughts of the last century[27]. Furthermore, in the four states with these highly unusual YFV outbreaks, we found that the outbreaks in each state coincided with peaks in the SPEI drought index over the last two decades. We also found that the outbreaks started in northern Minas Gerais which is the hottest and driest part of the southeastern region of Brazil. Our findings are further corroborated by a study conducted by Brazil’s National Center for Monitoring and Early Warning of Natural Disasters showing that the northern region of Minas Gerais was the area most severely affected by drought in 2016/2017 with an Integrated Drought Index at that time classified as extreme drought[27]. Our findings echo other studies which have also implicated extreme weather events as playing a role in triggering anomalous YFV outbreaks in Africa[28,29].

We postulate that environmental stress caused by drought decreased available habitat, forcing mosquitos and NHPs to congregate in high densities in the few remaining areas with adequate food and water. YFV is transmitted in Brazil by *Hemagogus* mosquitos which predominantly breed in pools of water in trees and feed on NHPs at the top of canopies, occasionally descending to bite humans when food is scarce. *Hemagogus* sp. can also travel long distances, up to 11.5 km in just a couple days [30]. During a drought, *Hemagogous* sp. could therefore seek out more optimal feeding and breeding grounds, concentrating around remaining water, NHPs, and humans. Howler monkeys frequently spend their days eating in the tops of canopies where *Hemagogous* sp. live [31]. Howler monkeys (*Alouatta sp*.) and marmosets (*Callithrix sp*.) are highly adaptable to fragmented, suboptimal habitats[31–33]. These NHPs also commonly cross between rural-urban boundaries and were the two predominant NHP genera identified in the epizootoic YFV surveillance program[1,2]. If NHPs and *Hemagogus* sp. were forced into higher density conditions in search of water and food, particularly near rural-urban borders, this could allow for amplification of YFV transmission once the hot rainy summer season started. Other studies have also shown that increased habitat stress and fragmentation promotes increased density of both *Hemagogus* sp. and Howler monkeys [6,34]. There is even some evidence that mosquitos may increase their biting rate during periods of drought due to dehydration, thereby further promoting viral transmission[35].

An additional factor that could have facilitated the spread of YFV from rural into urban areas at this time is open mosquito niches in urban areas. *Aedes aegypti* is typically the dominant mosquito species in urban centers in Brazil. However, severe outbreaks of dengue, chikungunya, and Zika virus in the years just preceding the yellow fever epidemic prompted aggressive mosquito control efforts. The *Aedes aegypti* index as measured in Rio de Janeiro had decreased 2 to 3 fold at the start of the YFV outbreak compared to the previous decade (Figure 5)[36]. Such a decrease in *Aedes aegypti* numbers could have allowed the typically forest dwelling *Hemagogus* sp. to take up residence in this open niche in urban centers. Although competition between *Aedes aegypti* and *Hemagogus* sp. has not been studied in this setting, such habitat competition between mosquitos has been documented in other studies[37,38]. This is further supported by several studies during this epidemic which found no evidence of YFV transmission by *Aedes aegypti*, despite them being competent vectors in lab studies, but rather transmission in urban areas predominantly by *Hemagogus* sp. and very rarely by other (non-aegypti) *Aedes* species [7,8].

**Figure 5.**
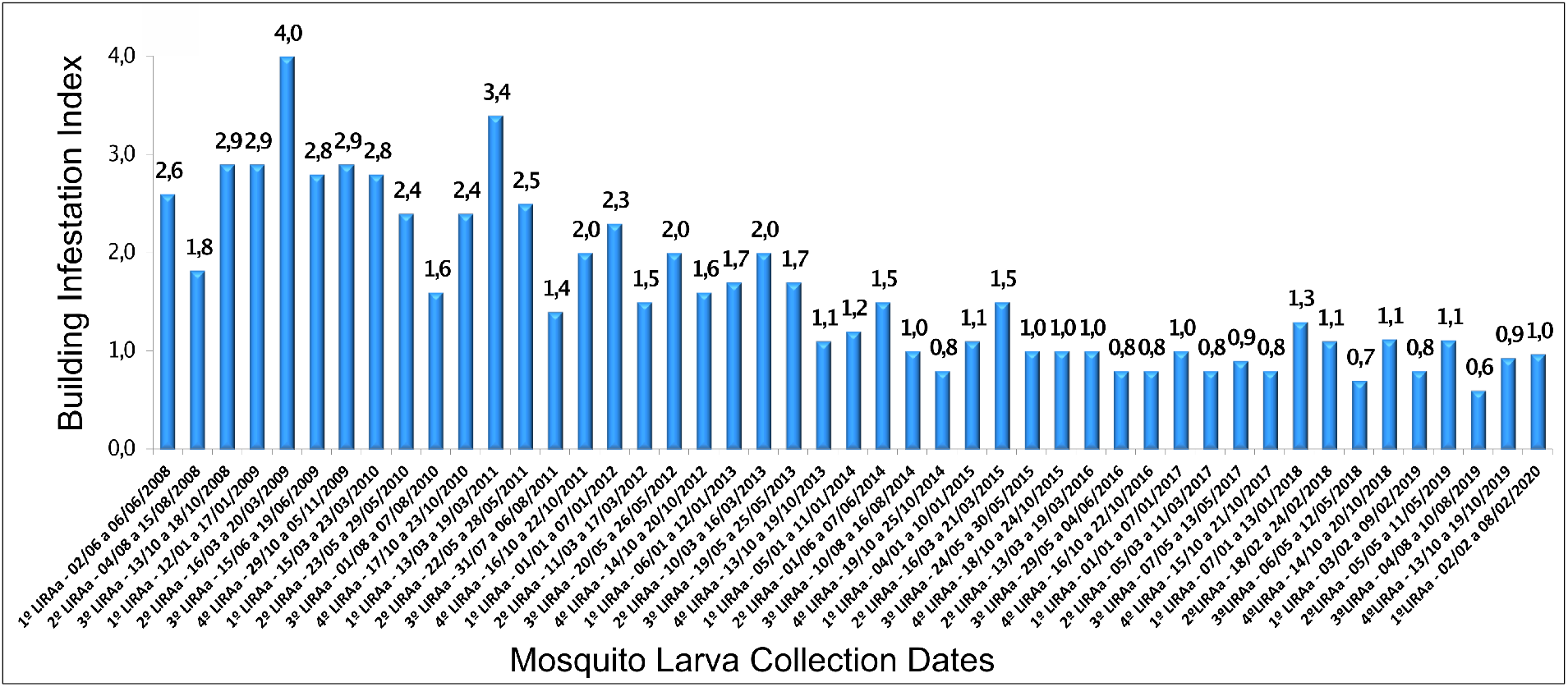
*Aedes aegypti* building infestation index in Rio de Janeiro, 2008-2020. Abbreviations: LIRAa = Levantamento de Índice Rápido para *Aedes aegypti*; English translation = Rapid Index Survey for *Aedes aegypti*. The LIRAa is the official measure for larval surveys of properties and provides an estimate of how many containers are infested with *Aedes aegypti* larva. The graph above is taken from the Brazilian Ministry of Health and is available at http://www.rio.rj.gov.br/web/sms/lira[36]. Mosquito larva surveys are conducted by the Ministry of Health in Rio De Janeiro 3-4 times a year over a 4-7 day period to measure the building infestation index for *Aedes aegypti* mosquito larva. The *Aedes aegypti* building infestation index has steadily decreased since 2008. When the yellow fever epidemic first hit Rio de Janeiro in 2017, *Aedes aegypti* numbers were at a ten year low. Low *Aedes* counts may be in part due to widespread *Aedes* mosquito control campaigns which were initiated due to outbreaks of dengue, chikungunya, and Zika in in the years just prior to and including 2016. This index is not available for other cities in southeastern Brazil, however, numbers were likely similarly low in other cities due to increased control efforts.

This study was limited by several factors, most notably by the episodic nature of this outbreak. The 2017-2018 yellow fever epidemic in southeastern Brazil was a single epidemic in a new geographic area, which makes it an important and interesting case study. However, without a longer historical record of epidemics in this region, it is difficult to ascertain whether spatiotemporal associations, such as the occurrence during a severe drought, are causal or merely coincidental. The seasonality of YFV also means that there are several months where yellow fever cases naturally subside. The absence of cases during those months in the middle of a drought period inherently decreases the model fit between case counts and drought index. To adjust for this, we conducted a secondary analysis limited to the summer months December to April during which YFV cases were present. The seasonality of YFV outbreaks during the hot, wet summer months coincides with peak summer travel. The strong association between travel and confirmed YFV cases could therefore be confounded by this seasonal travel effect given the the inherent overlap in these time periods. Despite this limitation, our hypothesis is supported by the fact that urban to rural travel was most strongly associated with confirmed YFV whereas travel from rural to urban centers did not have a significant association. This is also consistent with other data which supports an increased risk with rural / outdoor exposure.

This study was also limited by the data available from surveillance systems. The surveillance system primarily captured hospitalized cases of yellow fever and would have missed mild or asymptomatic cases. Surveillance and hospital systems are also not necessarily equally distributed throughout this region and were likely most robust in more urban areas, which could somewhat bias the distribution of cases to urban areas. The non-human primate cases were also captured using a convenience sampling of NHP carcasses that were tested for evidence of YFV infection; this sampling method would also likely bias towards a more urban distribution. Even with this potential urban bias in both human and NHP distribution, however, the epidemic clearly started in more rural areas and there was consistent evidence of increased risk with rural / outdoor exposure. Within the cases captured by the surveillance system, there was also some missing data. Although age and sex were nearly universally recorded, occupation was only available for about a third of confirmed cases. Additionally, this field was a write-in response which we sorted into four broad categories but detailed information about outdoor exposure for each occupation type was not available.

Despite these limitations, the spread of YFV into areas that had been YFV free for nearly a century signals changing conditions. Our study identifies patterns which suggest increased stress during a severe drought may have promoted the amplification of YFV transmission between mosquitos and NHPs at rural-urban boundaries, ultimately spilling over into a large human epidemic that spread from rural to urban areas. The reemergence of YFV in densely populated parts of Brazil raises serious concerns about the potential for reestablishing an urban transmission cycle which could involve *Aedes aegypti* in the future. More research is needed to understand what factors may have prevented YFV from reestablishing an infection cycle in *Aedes aegypti* during this epidemic and therefore what could be done to minimize that risk.

Lastly, the results of our study may have the potential to inform decision-making about YF immunization policies in southeastern Brazil. Currently in this region of the country, the official immunization calendar only recommends the YFV vaccine for children nine months and older who live in municipalities deemed to be at high risk of sylvatic YFV[39]. During the 2017-2018 outbreaks, the principal control strategy was ring vaccination of all age groups. The results of our study reinforce that sylvatic YF has the potential to spread rapidly across hundreds of kilometers among humans and NHPs in southeastern Brazil. In light of the potential for sylvatic YF epidemics to expand over considerable distances, decision makers might evaluate other immunization strategies including mass vaccination[40]. However, we acknowledge that the formulation of vaccination policy is a complex process that must take into consideration a variety of factors including the risk of adverse events, and the investment in infrastructure and training necessary to rollout mass vaccination.

